# Cortical adaptation to muscle fatigue does not alter early proprioceptive processing in primary sensorimotor cortex

**DOI:** 10.64898/2026.06.15.728001

**Authors:** Junru Chen, Toni Mujunen, Feiyue Li, Riku Nikander, Harri Piitulainen

## Abstract

Muscle fatigue potentially interferes with proprioceptive afference from peripheral “movement sensors”— the proprioceptors, which may hinder the crucial sensorimotor integration and thus locomotor performance. However, little is known about how muscle fatigue affects cortical processing of proprioceptive afference.

Twenty-four healthy volunteers (30.7 ± 6.5 yrs, 13 females) participated in the experiment, which included magnetoencephalography (MEG) recordings during ankle proprioceptive stimulation (2-Hz passive movements), and fatigue tasks comprised of isometric ankle plantar flexion. Corticokinematic coherence (CKC) between foot acceleration and MEG signals was examined before (PRE) and ∼3 min after (POST) the fatigue tasks to quantify the cortical proprioceptive processing.

CKC peaked in the gradiometer pairs above the foot region of the primary sensorimotor (SM1) cortex in each participant. CKC strength did not show significant difference between PRE and POST at 2 Hz (0.30 ± 0.12 vs. 0.30 ± 0.14, *p* = 0.981) or its first harmonic at 4 Hz (0.38 ± 0.14 vs. 0.37 ± 0.13, *p* = 0.724). However, 4-Hz MEG power was ∼30% lower in POST than in PRE. Surprisingly, fatigue-induced bilateral increase of alpha and beta power was observed in SM1 hand regions during the movement stimulation.

Our results indicated that the early processing of proprioceptive afference from the ankle joint was negligibly affected by muscle fatigue, or it recovered rapidly. The effects of muscle fatigue on the proprioceptive processing appear to extend beyond the primary somatotopic regions to bilateral SM1 neuronal networks. This cortical adaptation to muscle fatigue potentially preserves proprioceptive processing by modulating SM1 inhibitory neurons, offering a novel perspective for future research on proprioception.

## 1 Introduction

Muscle fatigue in the lower extremities primarily affects balance and locomotor performance in relation to proprioception (Helbostad et al., 2010; T. Li et al., 2024; Parijat & Lockhart, 2008). Proprioception encompasses various senses of body position and movement through afferent information arising from those mechanosensitive receptors called proprioceptors (Proske & Gandevia, 2012; Tuthill & Azim, 2018). It has been widely acknowledged that muscle fatigue can alter proprioceptive perception of the ankle joint, as evidenced by changes in the detection of motion (Lin et al., 2008), joint-position sense (Allen et al., 2010), and force sense (Brezovar et al., 2025; Proske & Allen, 2019). Due to the mismatch between perceived effort and actual output, it was suggested that central fatigue in the primary motor cortex affects proprioception by modulating transmission of the motor efferent copy to the somatosensory cortex (Smith et al., 2007; Søgaard et al., 2006; Taylor et al., 2006). However, there is no evidence on how the cortical processing of proprioception itself is altered by muscle fatigue.

Proprioceptors are highly sensitive to forces applied to the muscles, tendons, ligaments, and connective tissues surrounding the joints in a manner like cutaneous mechanoreceptors to touch. For example, the most sensitive ending of muscle spindles can detect below 10-μm changes in the muscle length (Brown et al., 1967; Roll et al., 1989). Golgi tendon organs on the other hand are activated with the rise of muscle tension (Fallon & Macefield, 2007; Jami, 1992). Various joint receptors adaptively respond to stretch acting on the joint capsule and firing tonically near the extremes of joint range of motion (V. G. Macefield, 2021; Rossi & Grigg, 1982). Proprioceptive afference, originating from the periphery, traverses the spinal cord towards the thalamic circuits, and subsequently reaches the cerebral cortex via the dorsal column-medial lemniscus pathway (Tuthill & Azim, 2018). The proprioceptive input project extensively to the somatosensory and motor cortices, with the most dense input to the primary sensorimotor (SM1) cortex contralateral to the peripheral proprioceptors (Ciccarelli et al., 2005).

Acute muscle fatigue can alter firing properties of the proprioceptors. In cats, firing rate of spindle Ia afferents is increased (Ljubisavljević & Anastasijević, 1994) and their sensitivity is reduced (Pedersen et al., 1998) in consequence of muscle fatigue. On the contrary, in human experiments, the spindle firing is reduced in fatigued muscles (G. Macefield et al., 1991). As substantiated by indirect evidence, an inhibitory spinal reflex—driven by the activation of group III and IV afferents in response to metabolic nociception and mechanical pressure (Taylor et al., 2016)—may lead to reduced γ-motoneuron excitability and thus less sensitive spindles (Avela et al., 1999; Bigland-Ritchie et al., 1986; Woods et al., 1987). Moreover, post-exercise modulation of cortical excitability and possible motoneuron excitability is partly mediated by residual afferent firing, such as group Ib and II afferents that are sensitive to intramuscular tension and static fiber length, in relaxed muscles (Gruet et al., 2013).

Furthermore, the fatigue-related changes in proprioception also occur in the brain. It has been reported that beta and gamma cortex-muscle coupling in ascending direction increased in the presence of isometric grip fatigue (Bedard et al., 2025; Liang et al., 2021). In numerous post-exercise electroencephalography (EEG) studies, increased alpha and beta activity was observed in the bilateral parietal cortex (Hosang et al., 2024), as well as magnetoencephalography (MEG) evidence from upper-limb fatigue studies (Fry et al., 2017; Tecchio et al., 2006), but this exercise-induced modulation was suspected to be overall brain oscillations that vanished immediately after the cessation of the exercise (Ciria et al., 2018). As is well known, the sensorimotor rhythms that are recorded over the Rolandic regions, typically 8–40 Hz in frequency, can be modulated by limb movements and tactile stimulation (Hari & Salmelin, 1997). For example, active movements with varying duration modulate sensorimotor rhythms to different degrees (Cassim et al., 2000; Pakenham et al., 2020), and even brief passive movement is robust modulator of ∼20-Hz rhythm (Parkkonen et al., 2015). Cortical responses to passive movement stimuli mainly reflect proprioceptive afference since they remain observable even in the absence of cutaneous feedback (Abbruzzese et al., 1985; Mima et al., 1996). However, the effect of muscle fatigue on cortical proprioception during passive movement have never been reported, although previous studies have revealed age-related inefficiencies of cortical proprioceptive processing (Piitulainen, Seipäjärvi, et al., 2018; Walker et al., 2020).

The cortical processing of proprioceptive afference is non-invasively assessed through corticokinematic coherence (CKC), which refers to the synchronization between cortical neuronal activity—measured with MEG or EEG—and limb kinematics (Bourguignon et al., 2011, 2015; Piitulainen et al., 2013b, 2020). The strength of CKC typically peaks at the movement frequency and its harmonics. The strongest CKC occurs simultaneously with the most regular movement stimuli (Mujunen et al., 2021), and it can be amplified with more comprehensive stimulation of homologous extremities, e.g., several fingers or single finger (Hakonen et al., 2022). Furthermore, CKC can be reliably estimated based on any peripheral signal picking the rhythmicity of movement (Piitulainen et al., 2013a). Importantly, CKC responses to rhythmic proprioceptive stimuli are reproducible for both the upper (Piitulainen et al., 2020; Piitulainen, Illman, et al., 2018) and lower limbs (J. Chen et al., 2025). In the present study, passive movement was elicited to the ankle joint in a rhythmic manner as proprioceptive stimulation.

We aimed to investigate how muscle fatigue affects cortical processing of proprioceptive afference from the ankle joint, and whether SM1 cortical activity is regulated by fatigue. We respectively quantified the degree of cortical proprioceptive processing using CKC strength and sustained movement-evoked field (MEF) amplitude, as well as alpha and beta power in the SM1 cortex during the movement stimulation, before and after isometric fatigue protocol. Our hypothesis was that (1) early cortical processing of proprioception would be hindered, i.e., CKC and MEF would be stronger, given that muscle fatigue is often accompanied by disturbed proprioceptor functioning and worse locomotor performance, and (2) SM1 cortex alpha and beta band activity would be stronger following fatigue, suggesting upregulation of inhibitory control over the related neuronal networks.

## 2 Materials and Methods

### 2.1 Participants

Twenty-four healthy adults (age 30.7 ± 6.5 yrs, BMI 23.3 ± 3.6, mean ± SD; 13 females) with no known history of neurological or musculoskeletal disorders were recruited for the study. Eighteen participants were right-footed and six were mixed-footed based on the Waterloo Footedness Questionnaire (Grouios et al., 2009). Prior to their enrollment, participants were requested to sign a written informed consent after a necessary understanding of the procedure. The study protocol was approved by the Ethics Committee of the University of Jyväskylä (approval number 633/13.00.04.00/2024), and the experiments were carried out in accordance with the Declaration of Helsinki.

### 2.2 Experimental Procedure

The measurements were conducted at the MEG lab in the Centre for Interdisciplinary Brain Research, University of Jyväskylä, Jyväskylä, Finland. Participants were first familiarized with the MEG environment and experimental devices. Then, a warm-up consisting of four 30-s in-place hopping was performed, with 30-s rest intervals in between. A custom-made MEG-compatible force device comprising a load transducer was used to perform fatigue tasks and measure ankle plantar force. The right foot was adjusted to their preferred position where the ankle joint was maintained ∼100° throughout the subsequent isometric contractions. The isometric maximal voluntary contraction (MVC) force was defined as the highest value among three utmost attempts, each lasting for ∼3 s. Figure 1 illustrates the experimental setup and procedures.

**Figure 1.**
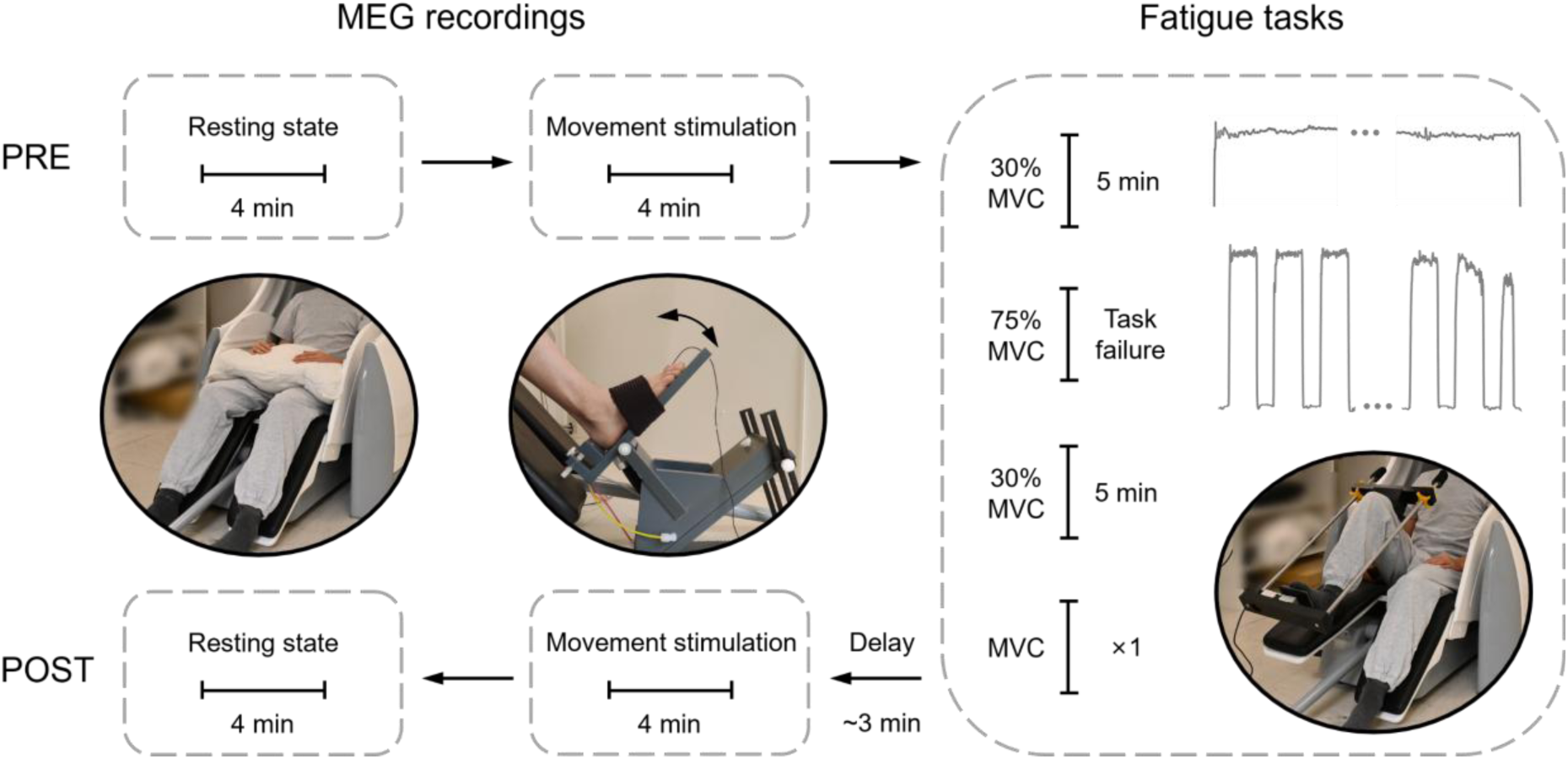
Experimental protocol and the device setup. For the resting state, participants were sitting comfortably in the MEG chair with their arms placing on the pillow in their laps, and fixated on a projected focal image on the screen. For the movement stimulation, participant’s right foot was fastened to the pneumatic actuator with elastic straps, otherwise identical to the resting. For the fatigue protocol, participants were instructed to adjust to their preferred position and maintain it throughout the tasks whenever possible. The force signals collected from a representative participant during steady and intermittent isometric contractions were respectively shown in the right panel. The delay means approximately 3-min recovery time after fatigue tasks.

#### 2.2.1 Fatigue Protocol

Muscle fatigue was induced through sustained submaximal isometric plantar flexion of the right ankle. Figure 1 right panel shows the fatigue protocol. The fatigue tasks consisted of two sustained 30% of MVC isometric contractions for 5 min each, and in between, a series of intermittent 75% of MVC isometric contractions. Participants performed 75% of MVC contractions 13 ± 4 (range 11–20) times, each lasting 20 s and separated by 10-s rests, until they failed to reach the target force. The target force levels were visualized using Signal software version 8 (Cambridge Electronic Design Ltd., Cambridge, UK) projected on a translucent screen placed 0.9 m in front of them. Participants were instructed to generate isometric force as steady as possible on a dotted horizontal line within ± 5% of MVC boundaries presented with solid lines. Upon the completion of the fatigue protocol, the force device was immediately replaced with the proprioceptive stimulator as quickly as possible, and the movement stimulation began ∼3 min (on average 228 ± 34 s) after MVC test.

#### 2.2.2 Proprioceptive Stimulation

Proprioceptive stimuli (i.e., ankle rotations) were generated using an ankle-movement actuator with pneumatic artificial muscles that are controlled by computer-generated trigger pulses (Piitulainen, Seipäjärvi, et al., 2018). The stimulation program was performed in Presentation software version 21.1 (Neurobehavioral Systems, Inc., Albany, CA, USA). Participant’s right foot was placed on the actuator, with the ankle secured in position using elastic straps. The movement actuator induced continuous passive dorsiflexion of the ankle joint (rotation range ∼5.5°, peak angular velocity 66°/s) at 2-Hz movement frequency. Proprioceptive stimuli were delivered to the ankle joint for 4 min, respectively, before (PRE) and after the fatigue tasks (POST). Participants were asked to fixate on a focal image through a rectangular hole in the cardboard in front of them. To minimize the auditory involvement of airflow noise from the actuator, we provided earplugs to participants and played 70 dB Brownian noise through flat panel speakers. In addition, resting-state MEG signals were recorded at the beginning and end of the MEG session for 4 min each.

### 2.3 Data Acquisition

The MEG recordings were carried out in a magnetically shielded room with a 306-channel Vectorview neuromagnetometer (Elekta Neuromag TRIUXTM, Elekta Oy, Helsinki, Finland) that contains 102 magnetometers and 204 planar gradiometers. MEG signals were bandpass-filtered online at 0.1–330 Hz and sampled at 1000 Hz. Foot kinematics was recorded using 3-axis accelerometers (ADXL335 iMEMS Accelerometer, Analog Devies Inc., Norwood, MA, USA) attached to the first proximal phalanx of the foot. Acceleration signals were lowpass-filtered at 330 Hz and sampled at 1000 Hz, time-locked to MEG signals.

The plantar force during fatigue tasks was measured with a rigid load cell (model YT-108, capacity 1962 N; Yongtai Measuring Instrument Co.,Ltd, Hangzhou, China) mounted on the custom-made device. Muscle activity was quantified using surface electromyography (EMG) recorded with Ag/AgCl electrodes (Ambu Neuroline 720, Ambu A/S, Ballerup, Denmark) taped on the tibialis anterior (TA), gastrocnemius medialis (GM) and soleus (SOL) muscles. The recording passband was 0.1–330 Hz for EMG signals and DC–330 Hz for force signal. EMG and force signals were then sampled at 1000 Hz, both time-locked with MEG signals.

### 2.4 Data Processing

#### 2.4.1 Force and EMG Signal Processing

Force signals were first band-pass filtered at 0.1–20 Hz with fourth-order Butterworth filter. Only the stable segments of force signals within the ± 5% target boundaries were included in further analyses. Force steadiness was quantified by the coefficient of variation (CoV) of force, calculated as the ratio of the mean to the standard deviation.

The raw EMG signals were time-aligned with the selected force segments. EMG signals were band-pass filtered at 5–295 Hz with fourth-order Butterworth filter. The root mean square (RMS) was then calculated using 250-ms sliding windows with 50% overlap. Median frequency (MDF) was estimated using Welch’s method with 1-s sliding windows and 50% overlap.

#### 2.4.2 MEG Signal Preprocessing

MEG raw data were first preprocessed using the temporal signal space separation (tSSS; Taulu & Simola, 2006) to suppress external interferences and correct for head movements in MaxFilter software version 3.0.17. The data was further preprocessed with MNE-Python software (Gramfort et al., 2013) to suppress physiological noise. After band-pass filtering the data to 1–40 Hz with a zero-phase finite impulse response filter (firwin in SciPy; FIR filter design using Hamming window), a fast independent component analysis (ICA; Hyvarinen, 1999) was implemented to decompose the data into 30 independent components. Then 2–4 ICA components corresponding to eye movements and cardiac activities were identified and ultimately discarded from the raw data.

#### 2.4.3 Coherence Analysis

Coherence analysis was carried out on the sensor level. Continuous MEG data were split into 2-s epochs with 80% overlap, leading to a frequency resolution of 0.5 Hz (Bortel & Sovka, 2007). Epochs with the signals exceeding 3 pT for magnetometers or 0.7 pT/cm for gradiometers were excluded to avoid contamination by any artifact not removed by the preprocessing. Next, coherence analysis (Halliday et al., 1995) was performed—yielding power-, cross- and coherence spectra, as well as correlograms—between all MEG sensors and the Euclidian norm of the three orthogonal accelerometer signals. The number of artifact-free epochs (560 ± 45, mean ± SD; range 483–595) was fixed equally for each participant between the two CKC recordings. The data from gradiometer pairs were combined in the direction of maximum coherence (Bourguignon et al., 2015), and then the gradiometer pair with the highest coherence value was selected from 10 gradiometer pairs over the left Rolandic foot region.

#### 2.4.4 MEG Steady-State Field Analysis

MEG signals were split into 1.5-s epochs and averaged with respect to the onsets of the movement stimulus. The resulting steady-state field was referred to as the sustained movement-evoked field (MEF). The sustained MEF was then bandpass-filtered with 1–45 Hz, and divided into plantar flexion and dorsiflexion phases according to the stimulus offsets. The peak-to-peak amplitude of MEF was calculated for the dorsiflexion phase, and finally extracted from the gradiometer pair that showed peak CKC at 2 Hz.

#### 2.4.5 Power Spectral Analysis of MEG Signals

Power spectral density (PSD) for the movement stimulation was computed concurrently with the coherence analysis. The resulting power spectra was then decomposed into the periodic and aperiodic component using FOOOF algorithm (Donoghue et al., 2020) with aperiodic mode “knee”, across the frequency range 1 to 49.5 Hz. Only the fitting results with *R^2^* > 0.90 were accepted for further analyses. After subtraction of the aperiodic component, the remaining PSD was used to calculate the power values at frequencies of interest: alpha band (8–13 Hz), beta band (15–25 Hz), movement frequency (2 Hz) and its first harmonic (4 Hz). In case the peaks of power spectra were lower than the fit, i.e., the periodic component at a given frequency is negative, its power was set to zero. The power in alpha band was defined individually for each recording as the mean within ± 1 Hz around the frequency with peak power. Similarly, the beta-band power was the mean within ± 2 Hz around the frequency with peak power.

### 2.5 Statistical Analysis

#### 2.5.1 Statistical Significance of Coherence

The statistical significance of the individual coherence levels was assessed under the assumption that the Fourier coefficients were linearly independent from one epoch to the next at each frequency of interest, while accounting for the use of overlapping epochs (Bourguignon et al., 2011; Halliday et al., 1995). To correct for multiple comparisons, the significance threshold was defined as *α* = 0.05/(Nf × Ns), of which Nf = 2, being the number of tested frequency bins, and Ns = 10, the number of preselected gradiometer pairs.

#### 2.5.2 Statistical Differences between PRE and POST

The normality of paired data distribution was initially assessed using the Shapiro-Wilk test. When the assumption of normality was accepted, paired t-test was used to evaluate statistical differences between paired data before and after fatigue, with effect sizes calculated using *Cohen’s d*. The magnitude of *Cohen’s d* was interpreted as follows: small (0.2–0.5), medium (0.5–0.8), and large (≥0.8) (Cohen, 1988). For non-normally distributed data, the non-parametric Wilcoxon signed-rank test was employed. Statistical significance was considered for all results when two-tailed *p* < 0.05.

#### 2.5.3 Correlations between CKC, MEF and MEG Power

To further estimate the effect of MEG signal-to-noise ratio on CKC, correlation analysis was performed for CKC strength with the corresponding MEF amplitude and MEG power. Pearson correlation coefficient (*r*) was used for normally distributed data, whereas for non-normal distribution, Spearman correlation coefficient (*r_s_*) was employed.

## 3 Results

All participants included in the final analysis (n = 24) showed significant CKC strength at the movement frequency (2 Hz) and its first harmonic (4 Hz). The 1/f-like component fit of MEG power spectrum was achieved in 22/24 participants during movement stimulation, with the remaining two excluded for poor model fit (*R^2^* < 0.90). Thus 22 participants (age 30.5 ± 6.7 yrs, 13 females) were included in the power spectral analysis.

### 3.1 The Degree of Muscle Fatigue

Non-fatigued and fatigued conditions were compared using the first 30 s of the first steady isometric contraction and the last 30 s of the second contraction. Figure 2 illustrates the effect of muscle fatigue on force steadiness and muscle activity. As shown in Table 1, the CoV of force significantly increased at the end of fatigue protocol, indicating reduced force steadiness in fatigued condition. The RMS of EMG amplitude increased significantly in TA, GM, and SOL muscles when they were fatigued. For the MDF of EMG signals, significant reduction was observed in TA and SOL following fatigue, but no significant difference was shown in GM. The MVC force was significantly lower immediately after the fatigue protocol (812 ± 134 N) than before fatigue (1044 ± 140 N, *p* < 0.001).

**Figure 2.**
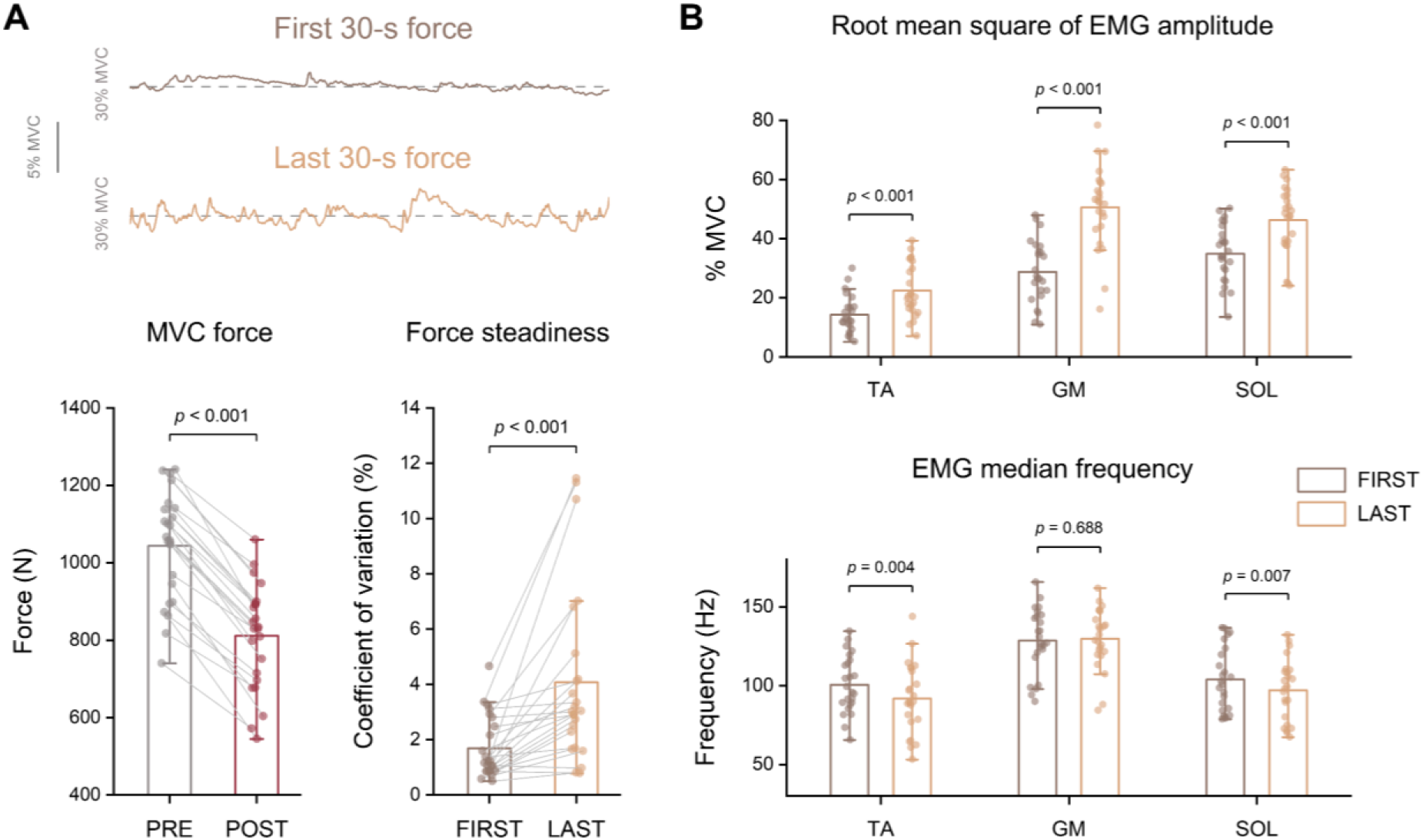
Force and EMG activity indicative of muscle fatigue. **(A)** Force fluctuations at the first and last 30 s of the fatigue protocol from a representative participant (upper panel). The force output of maximum voluntary contraction (MVC) before and after fatigue, and the coefficient of variation of steady isometric force (lower panel). **(B)** The root mean square of EMG amplitude in the tibialis anterior (TA), gastrocnemius medialis (GM) and soleus (SOL) muscles, and their EMG median frequency at the first and last 30 s of the fatigue protocol.

**Table 1.** Force steadiness and EMG activity.

|  |  | FIRST | LAST | *p* value |
| --- | --- | --- | --- | --- |
| Force CoV, % | | $1.69 \pm 1.12$ | $4.07 \pm 3.17$ | **< 0.001** |
| RMS, %MVC | TA | $14.2 \pm 6.3$ | $22.4 \pm 8.9$ | **< 0.001** |
| | GM | $28.8 \pm 10.7$ | $50.6 \pm 14.1$ | **< 0.001** |
| | SOL | $34.9 \pm 9.8$ | $46.2 \pm 11.2$ | **< 0.001** |
| MDF, Hz | TA | $100.6 \pm 18.2$ | $91.8 \pm 22.3$ | **0.004** |
| | GM | $128.5 \pm 20.6$ | $129.7 \pm 19.0$ | 0.688 |
| | SOL | $103.8 \pm 20.1$ | $97.1 \pm 19.5$ | **0.007** |

Note: Values are mean ± SD. CoV, coefficient of variation. MVC, maximal voluntary contraction. RMS, root mean of square. MDF, median frequency. TA, tibialis anterior. GM, gastrocnemius medialis. SOL, soleus.

### 3.2 Effect of Muscle Fatigue on CKC

Figure 3 presents the CKC strength from the gradiometer pairs showing the peak value at 2 Hz and 4 Hz. The CKC strength did not significantly differ between PRE and POST, at either 2 Hz or 4 Hz (Table 2). The grand average topographies showed, as expected, that CKC peaked consistently on the gradiometer pairs over the Rolandic foot area. In 10/24 participants, 2-Hz and 4-Hz CKC peaked in a different sensor, but the sensors were always the adjacent ones. The peak CKC sensor was largely consistent from PRE to POST recordings, with only 2/24 participants showing a shift to the adjacent sensor. One participant showed non-significant 2-Hz CKC of 0.04 in POST, while the corresponding value in PRE was 0.15.

**Figure 3.**
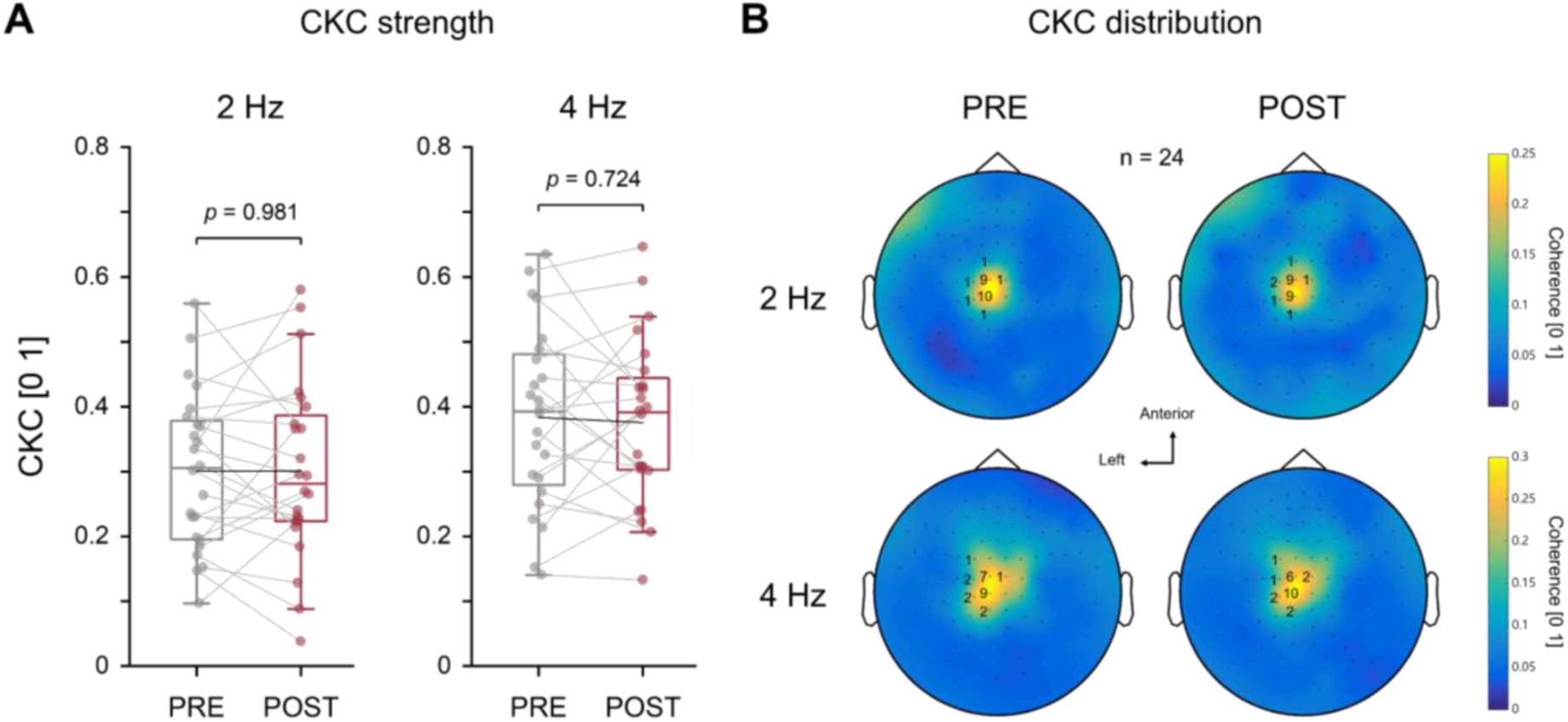
CKC strength and its topographic distributions (n = 24) before (PRE) and after fatigue (POST). **(A)** Peak CKC at 2 Hz and 4 Hz. Solid black lines connect the group mean values. The horizontal edges of the box indicate the interquartile range of the distribution, and the horizontal line in the box indicates the median. **(B)** The grand average topographies of 2-Hz CKC (upper panel) and 4-Hz CKC (lower panel). The number of peak CKC gradiometer pairs is marked in black.

**Table 2.**
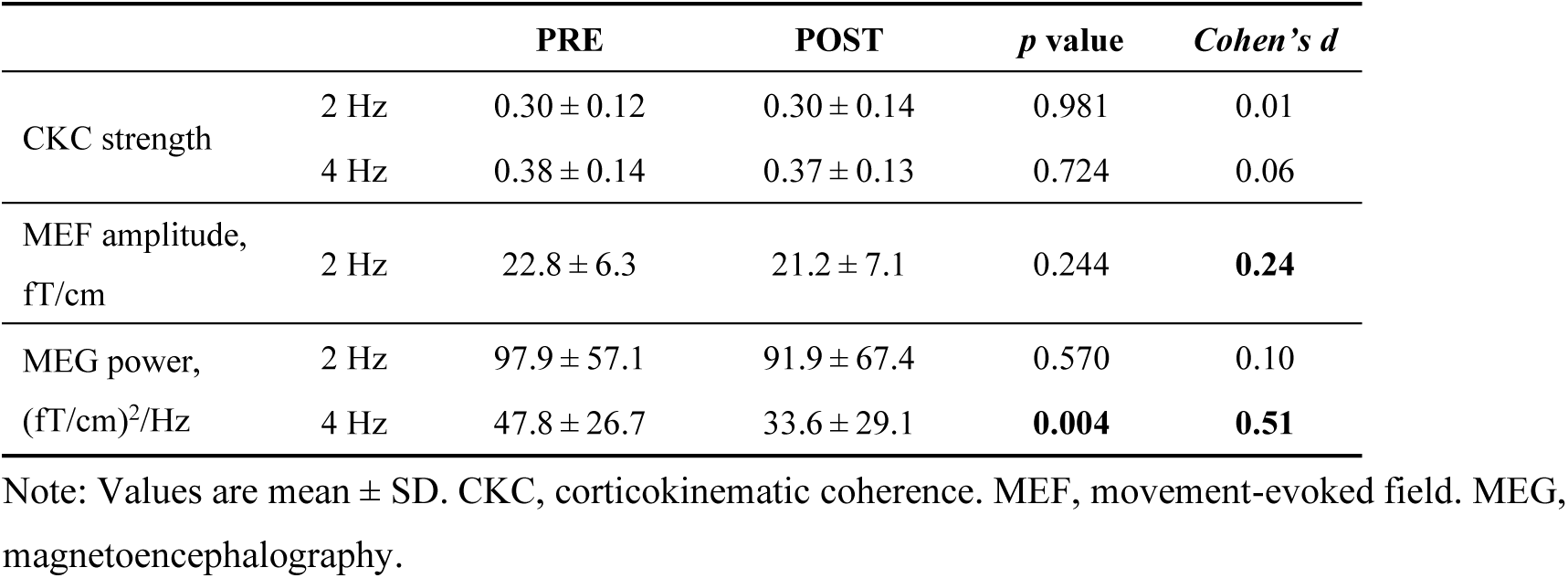
Cortical responses to proprioceptive stimulation.

### 3.3 Effect of Muscle Fatigue on Sustained MEF

Figure 4A shows sustained MEF that extracted from the gradiometer pairs showing strongest 2-Hz CKC. The sustained MEF amplitude did not significantly differ between PRE and POST (Table 2). Figure 4B illustrates the scatter plots and correlation between CKC strength and MEF amplitude. In PRE recording, stronger sustained MEF was associated with stronger CKC. However, in POST recording, the above correlative relation was no longer statistically significant. It is worth noting that the participant with the lowest CKC in POST unexpectedly had the highest MEF amplitude of 40.9 fT/cm.

**Figure 4.**
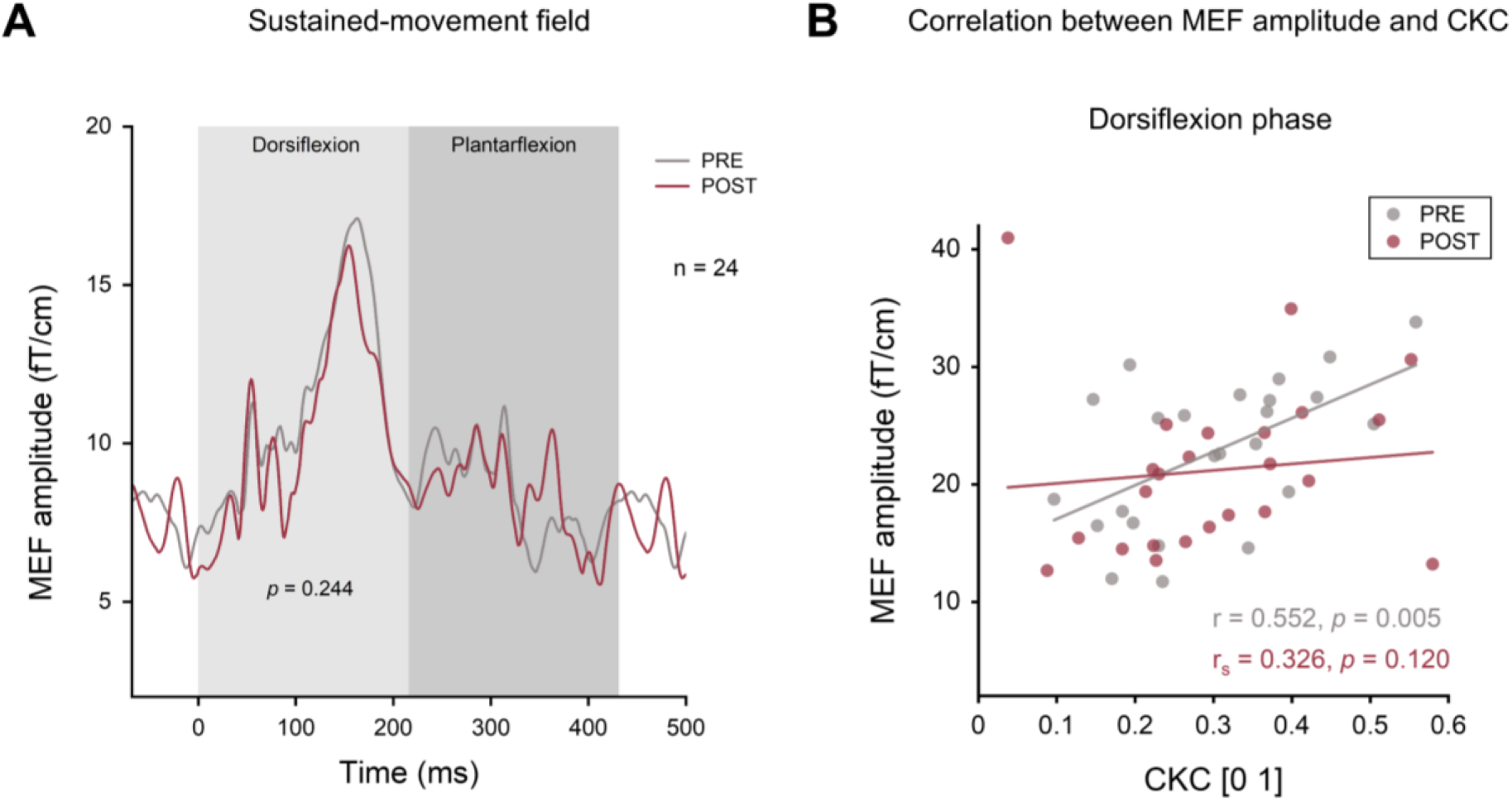
Sustained movement-evoked field (MEF) before (PRE) and after fatigue (POST). **(A)** Grand averages of MEF (n = 24) from the peak 2-Hz CKC sensors. The dorsiflexion and plantar flexion movement phases are highlighted with light and dark grey, respectively. The onset of movement stimulus is the beginning of dorsiflexion phase. **(B)** Correlation between 2-Hz CKC strength and the corresponding MEF amplitude.

### 3.4 MEG Power Modulation to Muscle Fatigue in Peak CKC Sensors

Figure 5 presents MEG power spectra extracted from the peak CKC sensors at 2 Hz and 4 Hz. MEG power at 2 Hz showed no significant difference between PRE and POST, while MEG power at 4 Hz was significantly weaker in POST compared to PRE (Table 2). CKC strength showed a significant correlation with both 2-Hz and 4-Hz power, both in PRE and POST recordings.

**Figure 5.**
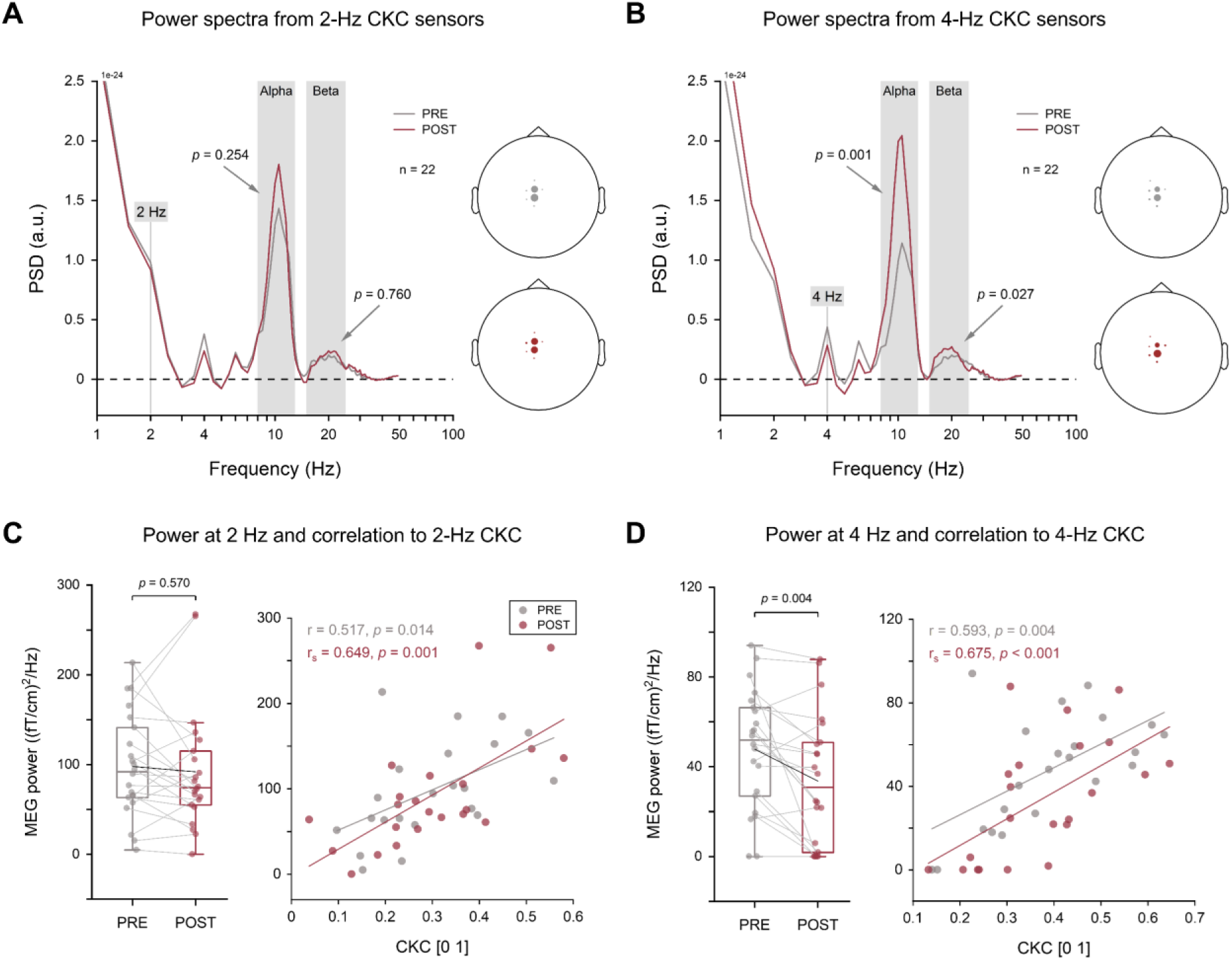
MEG power spectra extracted from the peak CKC sensors before (PRE) and after fatigue (POST). **(A)** Grand averages of power spectra (n = 22) from 2-Hz CKC sensors. The alpha (8–13 Hz) and beta bands (15–25 Hz) are highlighted with grey bars. The location of the sensors is marked with dots, with a larger dot indicating a greater number of sensors here. **(B)** Grand averages of power spectra (n = 22) from 4-Hz CKC sensors. **(C)** MEG power at 2 Hz and its correlation to 2-Hz CKC. **(D)** MEG power at 4 Hz and its correlation to 4-Hz CKC.

In addition, the power in alpha (8–13 Hz) and beta band (15–25 Hz) was also obtained from peak CKC sensors. For 2-Hz CKC sensors, no significant differences were observed between PRE and POST, in either alpha or beta power. However, for 4-Hz CKC sensors, both alpha and beta power were significantly stronger in POST than in PRE.

### 3.5 MEG Power Modulation to Muscle Fatigue in Peak Alpha and Beta Sensors

As shown in Figure 6, the topographic distributions of alpha- (8–13 Hz) and beta-band (15–25 Hz) mean power were plotted across all gradiometer pairs, as well as their differences between PRE and POST. Somewhat surprisingly, the most prominent lower-limb fatigue-induced changes in MEG power were observed in the sensors above the SM1 hand regions for both alpha and beta bands. To characterize these findings in more detail, we conducted exploratory analyses by selecting four gradiometer pairs with the largest power difference above the SM1 cortex contralateral and ipsilateral to the movement, respectively, and then averaged their power spectra. Based on exploratory findings, average alpha power was significantly stronger in POST compared to PRE in both contralateral and ipsilateral hand regions. For average beta power, the ipsilateral hand region was significantly stronger in POST than in PRE, however, the contralateral was not.

**Figure 6.**
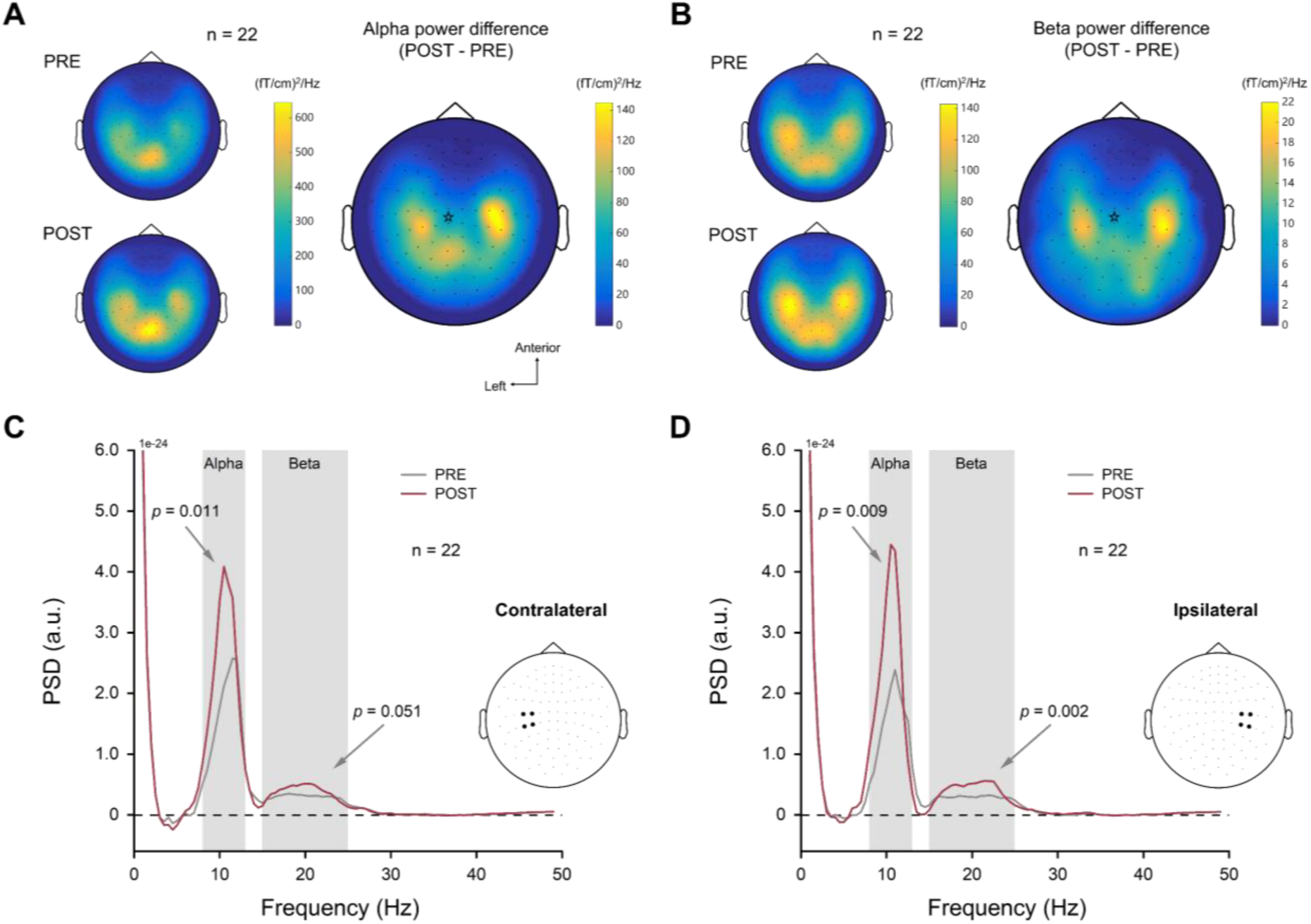
Grand average topographies of alpha (8–13 Hz) and beta (15–25 Hz) power during movement stimulation (n = 22) and the grand averages of power spectra extracted from the peak alpha and beta sensors before (PRE) and after fatigue (POST). **(A)** The topographies of alpha power and their difference between PRE and POST across all gradiometer pairs. The PRE values were always subtracted from the POST. The CKC node is shown as a star. **(B)** The topographies of beta power and their difference between PRE and POST across all gradiometer pairs. **(C)** Grand averages of average power spectra from four contralateral (left) gradiometer pairs. **(D)** Grand averages of average power spectra from four ipsilateral (right) gradiometer pairs.

## 4 Discussion

Our fatigue protocol induced clear muscle fatigue in plantar flexor muscles (GM and SOL). The proprioceptive stimulation resulted as clear cortical responses to the ankle movements. Contrary to our hypothesis, the early processing of cortical proprioceptive afference, i.e., CKC strength and MEF amplitude, was not significantly affected by muscle fatigue, only MEG power at first harmonic (4 Hz) of the stimulation frequency was reduced by the fatigue. However, we observed that MEG power in both alpha and beta band was significantly increased by muscle fatigue within a multi-node SM1 neuronal network, which may potentially affect the wide spread proprioceptive processing and sensorimotor integration in the SM1 cortex.

### 4.1 Muscle Fatigue Negligibly Affects Cortical Proprioceptive Processing

As expected, the evoked passive ankle rotations led to strong CKC with underlying sources located in the contralateral SM1 somatotopic foot region (J. Chen et al., 2025; Giangrande et al., 2024; Piitulainen et al., 2015; Piitulainen, Seipäjärvi, et al., 2018). Even when the muscles are relaxed, spindle Ia afferents encode the changes in fascicle length (G. Macefield & Knellwolf, 2018) and muscle force response to stretch (Blum et al., 2017). It has been shown that muscle fatigue can cause acute adaptation in the muscle spindle reactivity to passive stretch and vibration, such as increasing its firing rate and shortening stretch response latency (Nelson & Hutton, 1985). However, these fatigue-related changes in peripheral proprioceptors may not significantly impair the early cortical proprioceptive processing, since we observed no significant changes in CKC following muscle fatigue, nor the MEF amplitude. However, it cannot be ruled out that proprioceptive processing is unaffected by muscle fatigue, but with few minutes of recovery, its degree in the cortical neuronal networks appears negligible. It has been shown that the intensity of proprioceptive stimuli, e.g., larger vs. smaller movement range (Nurmi et al., 2023), do not significantly affect CKC strength, provided that the regularity of stimulation sequence is maintained (Mujunen et al., 2021). This highlights the “invariance” of early proprioceptive processing, indicating that the amount of proprioceptive afference reaching the cortex is finite and only needs to reach the threshold for triggering cortical responses (Mima et al., 1996; Seiss et al., 2002). Furthermore, applying sufficient vibration or electrical stimulation to fatigued muscles can effectively activate large-diameter afferents like group Ia and II (Avela et al., 1999; Griffin et al., 2001; Nelson & Hutton, 1985), which demonstrates that axonal conduction in these somatosensory afferents is not substantially impaired by metabolic exhaustion. On the other hand, the accompanying temperature elevation may alleviate the fatigue effects by increasing conduction velocity and metabolic turnover (Gray et al., 2006).

CKC is theoretically interpreted as a frequency-dependent measure of phase coupling between limb kinematics and cortical activity (Bourguignon et al., 2019). In fact, MEG signals mainly consist of inherent sustained electrophysiological “brain noise”, but also some external noise. Simulations have shown that improved signal-to-noise ratio may enhance the coherence strength in noisy signals (Muthukumaraswamy & Singh, 2011). Since CKC strength has often showed positive correlation with sustained MEF amplitude (J. Chen et al., 2025; Mujunen et al., 2021; Piitulainen, Illman, et al., 2018), amplitude coupling also plays an important role in the coherence estimation, especially in noisy signals like MEG. As expected, we observed significant positive correlation between CKC strength and MEF amplitude, but it was significant only before muscle fatigue, not after. Although MEF amplitude was not significantly altered following muscle fatigue, a small effect size between PRE and POST suggested that MEF changes (to be lower) did exist. In addition, 2-Hz and 4-Hz MEG power was consistently positively correlated with CKC strength both before and after fatigue. It is noteworthy that, as 4-Hz MEG power was reduced significantly, CKC would have been expected to decrease as well, in accordance with associations between MEG signal-to-noise ratio and coherence value. These indications may point to a “relative” increase in CKC strength, yet fatigued-induced elevation in brain noise has led to poorer coherence estimation.

### 4.2 SM1 Cortical Power is Regionally Modulated by Muscle Fatigue

We found that MEG power at 4 Hz was reduced by ∼30% following muscle fatigue, which suggests smaller neuronal population involved in the proprioceptive processing. Interestingly, in the same 4-Hz peak CKC sensor, alpha and beta power increased significantly by muscle fatigue, but not in 2-Hz peak CKC sensor. Since enhanced alpha and beta oscillations reflect “functional inhibition” in the cortex (Barone & Rossiter, 2021; Jensen & Mazaheri, 2010), the above findings could be interpreted as a selective inhibition of neurons dedicated to the movement harmonics. In other words, in the SM1 foot area, the cortex prioritizes the robust tracking of the fundamental movement rhythm over the processing of its finer nonlinear features.

Lower limb proprioceptive stimulation has been shown to activate the SM1 cortex beyond the expected foot region (F. Li et al., 2026; Mujunen et al., 2025). Our further power analyses revealed the most prominent fatigue-related increase of alpha and beta power was lateral to the foot somatotopy, peaking in the sensors above bilateral hand regions of the SM1 cortex. These findings suggest that fatigue-induced power modulations are not confined to a particular effector region but affects SM1 cortex dynamics on a wider basis. One explanation might be that the Rolandic foot area exhibits inherently weaker MEG sensitivity compared to the hand (Hillebrand & Barnes, 2002), as its cortical representation within the medial wall of the paracentral lobule is anatomically smaller and situated both deeper and more medially (Ciccarelli et al., 2005; Dobkin et al., 2004; Francis et al., 2009). Importantly, it has recently been suggested that, in addition to effector-specific regions for isolating fine action, inter-effector regions exist to integrate information from body movements (Gordon et al., 2023). In the present study, the alpha and beta power changes were qualitatively observed with bilateral peaks in the sensors above the inter-effector regions. Physiologically, this region-specific power modulation might relate to inhibitory neurotransmitter GABA concentration, as evidenced by the increased GABA concentration in upper-limb regions of the sensorimotor cortex after lower-limb exercise (Coxon et al., 2018). Thus, these exploratory findings should be explored in more detail in future studies.

Our results provide support for the view that alpha and beta power modulates inhibitory control over local neural populations (Z. Chen et al., 2026; Jensen & Mazaheri, 2010). We propose that muscle fatigue strengthens localized “inhibitory stencils” to constrain cortical processing in non-effector regions. This allows the processing of proprioceptive information to be more concentrated in the effector region, i.e., CKC node, in which alpha and beta activity is relatively weak and stable. In addition, the beta power appeared to be increased bilaterally in SM1 cortex following the muscle fatigue, consistent with the resting-state results from a grip fatigue study (Matta et al., 2024). As proposed by the “status quo” hypothesis, beta activity reflects the maintenance of the current sensorimotor state, which means that higher beta power results in greater difficulty in releasing cortical inhibition to initiate movement (Barone & Rossiter, 2021; Engel & Fries, 2010). Specifically, the motor cortex is less active once the peripheral proprioceptors become exhausted, or when the SM1 cortex itself is depleted as well. Indeed, upregulation of beta power has been linked to impaired function, such as less motor flexibility (Pierrieau et al., 2025). From the perspective of proprioceptive processing, stronger SM1 beta power is closely associated with early-stage diabetic neuropathy (Mujunen et al., 2025). Furthermore, impaired movement execution observed in aging (Rossiter et al., 2014) and Parkinson’s disease (Little & Brown, 2014) is directly related to higher beta power at rest. Taken together, the power modulation of cortical responses to muscle fatigue, as a form of “functional inhibition”, may be regionally specific as opposed to a global phenomenon.

### 4.3 Limitations and Future Perspectives

In comparison with proprioceptive stimulation with a long interstimulus interval (ISI ∼3 s; Smeds et al., 2017), acute exercise-induced fatigue is more capable of rapidly collecting sufficient stimuli for higher signal-noise-ratio. It is noteworthy that rapid rhythmic stimulation may mask some fatigue-related effects on proprioceptive signals, since the temporal expectation of regularities can boost the neural processing of anticipated movements (Nobre & Van Ede, 2018; Rohenkohl et al., 2012), and even result in time-locked cortical responses to omitted stimuli (Andersen & Lundqvist, 2019) given that the interstimulus interval is stable.

Movement-related artifacts derived from rhythmic ankle-joint rotations were sometimes observed at the lowermost MEG helmet sensors (in 6/24 participants), and were similarly reported in prior lower-limb CKC studies (J. Chen et al., 2025). Nevertheless, the presence of distinct CKC peaks at the vertex sensors located above the SM1 cortex suggests that the artifacts did not mask the actual neurophysiological responses. It appears that continuous lower-limb movement stimuli may produce unintended “oscillating” head movements when compared to the intermittent ones (Mujunen et al., 2022). If necessary, individualized MRI-based head casts could be used to reduce fluctuations of the head position (Meyer et al., 2017). We also recommended to employ a moderate movement range and angular velocity for ankle rotation, in conjunction with precise alignment of the ankle joint and actuator rotation axes, which can effectively avoid any redundant movements.

Due to replacement of the proprioceptive stimulator, we could not avoid a few minutes of recovery period between the fatigue tasks and subsequent proprioceptive stimulation. During the recovery, muscle reperfusion may partly attenuate the effects of group III and IV afferent activations by decreasing the accumulation of metabolites (Amann et al., 2015; Carroll et al., 2017). It has been reported that, despite rapid recovery of MVC force within few minutes, resting twitch forces evoked by motor nerve stimulation do not recover appreciably within 20 min after sustained low-intensity contractions (Søgaard et al., 2006). We recommend that caution should remain when estimating the recovery time and extent of fatigue, given the task-dependent nature of fatigue (Enoka et al., 2011) and its dependence on the measures (Krüger et al., 2019). Future studies should pay attention to the fatigue recovery or directly track cortical proprioceptive processing as fatigue evolves.

## 5 Conclusion

The cortical processing of proprioceptive afference from the ankle joint was negligibly affected by muscle fatigue or recovered within minutes. Interestingly, fatigue-induced increase of alpha and beta power was observed, and surprisingly, beyond the somatotopic foot region in the bilateral SM1 hand regions. We interpreted this observation as region-specific cortical inhibition that may allow more focused lower limb proprioceptive processing compensating the effects of the muscle fatigue. Further studies should investigate the patterns of cortical proprioceptive processing with fatigue evolution and also their implication for patients with chronic pathological fatigue (Chaudhuri & Behan, 2004).

## Author Contributions

**Junru Chen**: Conceptualization, Methodology, Software, Validation, Formal analysis, Investigation, Data curation, Writing – original draft, Visualization, Project administration. **Toni Mujunen**: Methodology, Software, Writing – review & editing, Data Curation. **Feiyue Li**: Investigation, Writing – review & editing. **Riku Nikander**: Writing – review & editing, Supervision. **Harri Piitulainen**: Conceptualization, Methodology, Resources, Writing – review & editing, Supervision, Project administration, Funding acquisition.

## Data Availability Statement

The data that support the findings of this study are available from the corresponding author upon reasonable request.

## Abbreviations

CKC: corticokinematic coherence
MEF: movement-evoked field
MEG: magnetoencephalography
SM1: primary sensorimotor cortex
MVC: maximal voluntary contraction
EMG: electromyography
TA: tibialis anterior
GM: gastrocnemius medialis
SOL: soleus
CoV: coefficient of variation
RMS: root mean of square
MDF: median frequency
PSD: power spectral density.

## ACKNOWLEDGMENTS

This work was supported by grants from the Research Council of Finland (#296240, #326988, #327288 and #361732) to HP, and “Brain changes across the life-span” profiling funding (#311877) to University of Jyväskylä. This work was also supported by a PhD scholarship from the China Scholarship Council (#202306060030) to JC.

The authors sincerely acknowledge the technical supports from Markus Kerminen for hardware manufacturing and Sakari Vekki for controller programming, and Heidi Pesonen for her valuable assistance with data collection, and Santtu Seipäjärvi for his insightful comments on the study findings. In addition, the authors would like to thank all the volunteers who participated in the experiment.

